# Acoustic Salience Drives Pupillary Dynamics in an Interrupted, Reverberant Task

**DOI:** 10.64898/2026.03.31.715639

**Authors:** Victoria I. Figarola, Wusheng Liang, Sahil Luthra, Eric W. Parker, Matthew B. Winn, Christopher A. Brown, Barbara Shinn-Cunningham

## Abstract

Listeners face many challenges when trying to maintain attention to a target source in everyday settings; for instance, reverberation distorts acoustic cues and interruptions capture attention. However, little is known about how these challenges affect the ability to maintain selective attention. Here, we measured syllable recall accuracy and pupil dilation during a spatial selective attention task that was sometimes disrupted. Participants heard two competing, temporally interleaved syllable streams presented in pseudo-anechoic or reverberant environments. On randomly selected trials, a sudden interruption occurred mid-sequence. Compared to anechoic trials, reverberant performance was worse overall, and the interrupter disrupted performance. In uninterrupted trials, reverberation reduced peak pupil dilation both when it was consistent across all stimuli in a block and when it was randomized trial to trial, suggesting temporal smearing reduced clarity of the scene and the salience of events in the ongoing streams. Pupil dilations in response to interruptions indicated perceptual salience was strong across reverberant and anechoic conditions. Specifically, baseline pupil size before trials did not vary across room conditions, and mixing or blocking of trials (altering stimulus expectations) had no impact on pupillary responses. Together, these findings highlight that stimulus salience drives cognitive load more strongly than does task performance.

## I. INTRODUCTION

Laboratory studies of attention often utilize anechoic listening conditions that do not reflect the acoustic complexity of real-world acoustic settings. In naturalistic listening environments, such as classrooms, hospitals, and other common settings, listeners must focus on a target source while ignoring competing sounds and background noise. This ability to attend to different parts of an acoustic signal is critical for successful communication, yet it becomes more difficult when reverberation and unexpected distractions distort the target signal or distract a listener. In such conditions, the auditory system must support sound segregation (parsing the sound mixture to individual sound sources) and sound selection (prioritizing the target sound source for further processing) (Best et al., 2006; Middlebrooks and Waters, 2020; Shinn-Cunningham, 2005).

Reverberation occurs in common place environments, resulting in sound energy reflecting off surfaces and reaching the listeners’ ears at multiple delays (Bradley et al., 2003; Duquesnoy and Plomp, 1980; Guski, 1990; Nabelek et al., 1989; Soulodre et al., 1989). Though early reflections can sometimes improve perception of a sound by adding energy without significantly degrading the content, they nonetheless degrade features important for source segregation, including spatial cues and pitch cues. As a result, even when the target stream remains audible, maintaining attention over time in natural, reverberant settings can require substantial cognitive resources. Later reflections not only make it harder to segregate sound sources, they smear acoustic energy over time, which can reduce information-conveying temporal modulation, increase background energy, and reduce the perceptual distinctiveness of individual sound sources (Culling et al., 2003; Culling and Stone, 2017; Duquesnoy and Plomp, 1980; Kidd et al., 2005; Lavandier and Culling, 2007; Lee and Shinn-Cunningham, 2008; Nabelek et al., 1989; Picou et al., 2016; Ruggles and Shinn-Cunningham, 2011). As a result, these larger-scale distortions from late reflections reduce speech intelligibility and speech perception in addition to impairing source segregation. Thus, in reverberant settings, maintaining selective attention to make sense of a target source in the presence of other competing sounds may be especially difficult.

In addition to ongoing reverberation distorting acoustics, everyday listening also often involves salient interruptions that can capture attention and disrupt ongoing processing. Such events are behaviorally relevant because they may trigger involuntary, bottom-up attention to the interrupter; interfere with encoding of target content; and require reorienting of top-down, selective attention back to the ongoing target stream (Liang et al., 2022; Mondor and Zatorre, 1995; Parmentier, 2014; Parmentier et al., 2011; Shapiro et al., 1997). A recent series of psychophysical experiments explored the behavioral impacts of interruption during spatial selective attention (Figarola et al., 2025; Liang et al., 2022, 2025). In these experiments, listeners were asked to selectively attend to a target syllable stream from one direction while ignoring a distractor syllable stream from a different direction. On randomly selected trials, an interrupter occurred, which disrupted recall of target syllables, especially the syllable immediately after the interruption. In anechoic simulations, varying the working memory and the temporal position of the interrupter did not change the basic pattern: An interrupter impaired listeners ability to recall syllables from the target stream, especially the syllable immediately after the interruption (Liang et al., 2022, 2025). One study that used target and distractor streams of five syllables each found that including naturalistic reverberation had little impact on the ability to recall a target stream for uninterrupted trials; however, in interrupted trials, the interrupter hurt performance more in reverberant than in anechoic simulations (Figarola et al., 2025).

Although behavioral measures can provide some insight into how reverberation and interruption affect cognitive processes, physiological measures can reveal perceptual and cognitive differences that may be unobservable from behavior alone. One such measure is pupil dilation. Pupil changes can be broken down into relatively slow tonic responses and more rapid phasic responses. Prior studies have shown that tonic responses are linked to arousal, alertness, and overall listening effort (Beatty, 1982; Bradley et al., 2008; Kahneman and Beatty, 1966; Vazey et al., 2018; Winn et al., 2018), and relate to performance accuracy (Heitz et al., 2008; McGinley et al., 2015; Winn et al., 2018). Anticipation of difficult trials can even elicit larger tonic pupil dilation before the stimulus is presented (McCloy et al., 2017). In contrast, rapid phasic responses relate to stimulus events and reflect surprise, momentary changes in cognitive effort, and even decision-making processes (Beatty, 1982; Dayan and Yu, 2006; Lisi et al., 2015; Soulodre et al., 1989; Vazey et al., 2018; Winn et al., 2018; Zekveld et al., 2018; Zhao et al., 2019b). Peak latency and amplitude in phasic responses correspond to the maximum processing time and processing load, respectively (Koelewijn et al., 2015; Zekveld et al., 2011). Pupillary dynamics thus capture continuous, time-resolved changes in cognitive and sensory processing that cannot be inferred from behavioral accuracy alone. This makes pupillometry well-suited for evaluating how processing evolves during sustained auditory attention and how listeners respond to salient disruptions embedded in ongoing streams.

Here, we measured behavioral performance and pupil dilation simultaneously while listeners performed a spatial attention task in which target and masker streams were presented either in anechoic or reverberant conditions. Our paradigm closely followed that used in the previous study investigating interactions between reverberation and interruption (Figarola et al., 2025), but used slightly shorter target / distractor streams to reduce trial duration (four vs. five syllables per stream). On a subset of trials, a brief interruption occurred during the ongoing stream. We considered two, not mutually exclusive, hypotheses: Reverberation could make the onsets of all events, including the interrupter, less salient by smearing the acoustic features of the ongoing streams; this would reduce phasic responses to syllables in the target and distractor streams as well as making the interrupter onset less distinct, and thus less disruptive. Additionally, reverberation could make the task harder overall, both by decreasing the clarity of the target speech and by smearing out the spatial cues allowing listeners to focus and reorient spatial attention, compounding the detrimental effect of the interrupter. Physiologically, we expected the tonic pupil response linked to effort to be greater in reverberation than in anechoic settings due to the increased difficulty; however, we expected the phasic pupil response to ongoing stimulus events (including the interrupter) to be reduced by reverberation, as the acoustic smearing of reverberation should reduce the salience of event onsets.

We found that reverberation impaired behavioral performance and altered pupil dynamics. Unlike in the previous study (Figarola et al., 2025), we found that performance was worse in reverberant than in anechoic trials, even for uninterrupted trials. Moreover, the current study found a greater effect of the interrupter in anechoic trials compared to reverberant trials, while the previous study found a smaller, but opposite effect. The baseline (tonic) pupil responses were equivalent across anechoic and reverberant settings, rather than being larger in reverberation as we hypothesized. In uninterrupted trials, reverberation reduced phasic responses, consistent with our expectations that onsets in the ongoing streams would be less distinct; however, interruptions elicited similar, robust pupil responses across the two environments, suggesting that onsets of new streams were equally salient in anechoic and reverberant settings. Together, these results demonstrate that the effect of reverberation on performance depends on task demands: there are larger behavioral effects when cognitive load is relatively low and acoustic differences across room conditions play a relatively greater role in determining performance. In addition, pupillometry reveals differences in how reverberation shapes underlying sensory and cognitive processes beyond what can be inferred from behavioral accuracy alone.

## II. MATERIALS AND METHODS

### A. Participants

We wished to test for a significant interaction between the effect of reverberation and the effect of interruption on pupil dilation. We therefore conducted a power analysis on pilot data from four participants (estimated effect size Cohen’s d = 0.37, α = 0.05, power = 0.8). This analysis determined that a minimum sample size of 59 participants would be required in each experiment.

Studies were approved by the Institutional Review Board of Carnegie Mellon University. Participants were recruited from the Carnegie Mellon University community and were compensated for their time in either cash or partial course credit. All participants were native English speakers, had normal or corrected-to-normal vision (no hard contact lenses), had no history of eye diseases or nystagmus, and did not have diabetes.

Participants were screened to ensure they had no diagnosed hearing impairment or neurological disorder affecting auditory function and had pure-tone audiometric thresholds within 20 dB of normal (125 Hz to 8000 Hz). Finally, all participants were screened to ensure they were able to perform the experimental task; those who did not achieve an 80% pass rate were excluded (see Training section below).

We recruited 99 human participants between the ages of 18 and 35. Of the 99 participants, 60 passed all screenings and the training task and completed the experiment (21 men, 39 females, 1 not reported; age 22.23 ± 4.02 years), 24 were excluded for not passing the hearing screening, six were excluded due to technical difficulties, and nine were excluded due to missing data (see Methods below).

A follow-up, small scale study was also performed to investigate whether differences in perceived difficulty between anechoic and reverberant blocks influenced results in the main experiment. We recruited 19 participants between the ages of 18 and 35. Of the 19 participants, 15 passed all screenings and the training task and completed the experiment (10 men, 5 females; age 19.53 ± 1.51 years), three were excluded for not passing the hearing screening, and one was excluded due to missing data (see Methods below). Results for this follow-up study are in the Supplementary Material (FIG S1 and S2).

### B. Stimuli

Both target and distractor streams comprised four syllables randomly drawn with replacement from the same set. The set consisted of three syllables, [/ba/, /da/, /ga/], produced by a male, native English-speaking talker. Each syllable had a duration of 450 ms. The target and distractor streams differed only in their location, with one stream spatialized to 30 degrees left and the other to 30 degrees right of center. Which stream was left and which right was chosen randomly and independently for each trial. Within each stream, syllables were separated by 150 ms of silence. The two streams were temporally interleaved, with the leading stream starting 500 ms after the cue ended and the lagging stream starting 300 ms later. Thus, the syllable-to-syllable onset time within each stream was 600 ms, but the two streams together created an isochronous mixture with syllables alternating between target and masker every 300 ms. The target stream started first in half the trials and the distractor was first in the other half; in half the trials, the first stream was on the left and in the other half, the first stream was on the right. In 50% of the trials (randomly selected), a novel “interrupter” (duration = 250 ms) occurred 125 ms before the onset of the 2nd syllable in the target stream, from 90 degrees left/right of center contralateral to the target.

Interrupters were brief, non-speech sounds selected to be acoustically and perceptually distinct from the syllable streams, ensuring they remained salient attention-capturing events even under reverberant conditions. A set of 80 unique (half of trials) interrupter sounds was drawn from the internet and included everyday events (e.g., cat meows, door slams, dog barks). Each interrupter was presented only once per participant.

### C. Spatialization

All syllables and interrupters were spatialized by convolving the monaural source with binaural impulse responses. Reverberant simulations were generated by convolving the sources with binaural room impulse responses (BRIRs), while pseudo-anechoic simulations were generated by convolving the sources with BRIRs that were time windowed so that they included only the direct path from source to the ears.

1. Pseudo-anechoic: the BRIRs measured in a classroom were obtained from an existing dataset recorded using a KEMAR in a classroom at Boston University (RT60 = 743 ms; (Shinn-Cunningham et al., 2005). Recordings were made from a central listening position with a loudspeaker located 1 m from the listener at 0-deg elevation, with the ears positioned approximately 1.5 m above the floor. Impulse responses were recorded at azimuths from 0° to 90° in 15° increments.
2. Reverberant BRIRs were recorded in a concert hall (RT60 = 1.919 sec). The BRIRs and the RT60 were calculated using a Python ‘psylab’ package (Brown, 2017). Specifically, the BRIRs were calculated by taking the Fourier transform of recorded sweep signals divided by the Fourier transform of the sweep; the RT60 was calculated based on the (Schroeder, 1965) method. Measurements were taken using a loudspeaker (MSP5A, Yamaha, Shizuoka, Japan) placed two meters away from the center of KEMARs head at 0-deg head elevation. The KEMAR had a microphone in each ear canal (Bruel & Kjaer; model 4134). A five-second-long sweep signal (50 Hz – 18 kHz) played from the loudspeaker and the response was measured at the KEMARs ear canal entrance. BRIRs were measured for azimuths 0° to 90° in 15° increments relative to the KEMAR. These azimuths were obtained by using a protractor and rotating the KEMAR head to the required angle relative to the loudspeaker. (All other equipment was kept the same).

### D. Eye-tracking procedure

Participants sat with their head fixed on a chinrest in front of a monitor (24-inch with resolution of 1920 × 1200 pixels and a refresh rate of 60 Hz) in a lit and sound-attenuated room. They were instructed to maintain fixation on a white cross centered with a black background on a computer screen. An infrared eye-tracking camera (EyeLink 1000 Desktop Mount, SR Research Ltd.) was placed below the monitor at a horizontal distance of 54 cm. Auditory stimuli were delivered through a Focusrite Scarlet Gen 3 sound card connected to a pair of headphones (Sennheiser brand). The loudness of the auditory stimuli was adjusted to a comfortable listening level for each participant. The standard 9-point calibration procedure for the EyeLink system was conducted prior to the experiment, as well as a 9-point validation procedure. During the experiment, the eye-tracker continuously monitored the participant’s gaze position and pupil diameter. Participants were instructed to blink naturally and rest in between blocks.

### E. Training

After eligibility screening, all participants were tested in a 5-minute training paradigm (implemented on https://gorilla.sc; (Anwyl-Irvine et al., 2020)). The training consisted of presenting one or more blocks of eight uninterrupted trials (4 anechoic and 4 reverberant) identical to uninterrupted trials in the main task (see below). Participants achieving 80% correct or better across a block of eight training trials progressed to the full study (Liang et al., 2022). Participants were given up to 5 chances to achieve the training criteria before being dismissed.

### F. Main Task

In all experiments, participants performed 160 trials in which they heard two competing, temporally interleaved, spatialized streams from opposite hemifields. At the start of each trial, a single spatialized /ba/ syllable played from either 30-degrees left or right, cueing the listener to attend to the stream at that location (the target) and ignore the other (the distractor). Once the stream ended, there was a 3-second silent interval to allow the pupils to return to baseline. Participants were then asked to report back the sequence of four target syllables played in the target stream via a computer keyboard. After their response, there was an additional three seconds of silence before the next trial began (FIG 1).

**FIG. 1.**
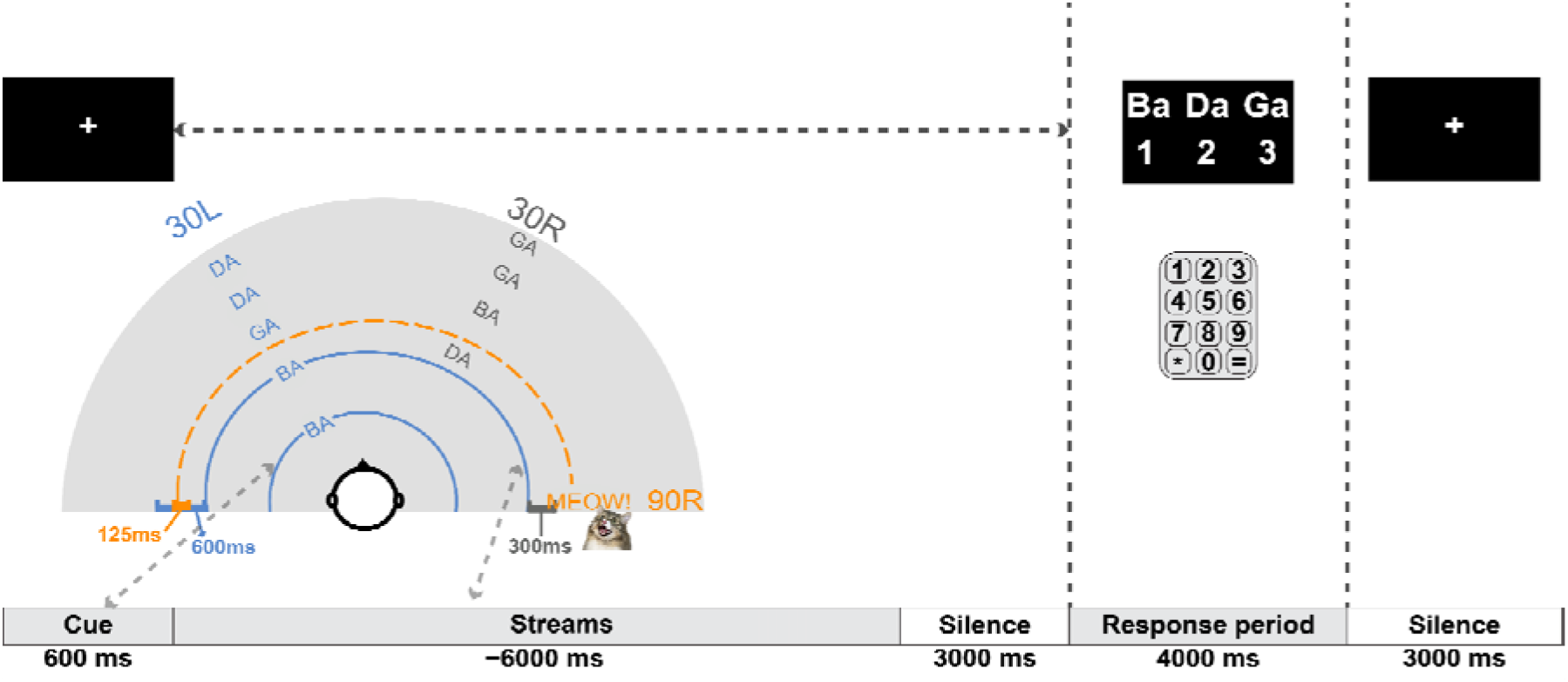
Schematic of Experimental Procedure. Each trial began with a visual fixation and an auditory cue indicating the target stream direction, followed by the presentation of the competing, temporally interdigitated target and distractor streams, played from +30 and -30 degrees. Following the presentation of the streams, there was a 3s silence, a response period in which subjects were asked to report the target syllables (in order), and a final 3s silent period. Subjects were randomly assigned to always hear stimuli with the left stream leading or the right stream leading. The cue defining the target direction was randomly chosen on each trial to be either left or right with equal likelihood; thus, the target stream was temporally leading on half the trials and lagging in the other half. The example shown in the top-down diagram represents a left-leading target stream (i.e., the cue appeared on the left and then the left stream began before the right stream).

The 160 trials were divided into eight blocks of 20 trials each, where the acoustic condition (anechoic or reverberant) was consistent across all stimuli within a block. Half of the blocks presented anechoic simulations and half reverberant simulations, alternating across blocks. Half of the participants began with an anechoic block and half with a reverberant block.

The direction of the leading stream was manipulated between subjects: half of the subjects always heard stimuli with the left stream leading and half always heard stimuli with the right stream leading. Importantly, all subjects were presented with an equal number of target-left and target-right trials. Although target stream direction and target timing (leading vs. lagging) were not counterbalanced within subjects, this design choice should not affect our primary question concerning the effect of the interrupter and its interaction with reverberation.

Within each block, trials were fully counterbalanced such that half the trials were uninterrupted and half contained an interrupter, and half had the target on the left and half the target on the right. Trial order was randomized independently for each block and each participant. Participants were instructed to rest in between blocks.

In the follow-up experiment, the same conditions were presented, but the blocking was changed. Specifically, each of the eight blocks either contained only uninterrupted or only interrupted trials, while anechoic and reverberant trials were randomly intermingled with each block.

### G. Behavioral Analysis

Behavioral performance was computed as the proportion of target stream syllables correctly recalled, averaged across trials within each participant and condition (anechoic/reverberant x uninterrupted/interrupted). Proportion correct performance was analyzed across three factors: syllable position (1-4), interrupted condition (interrupted vs. uninterrupted), and room condition (anechoic vs reverberant). A 2 (Interruption) x 2 (room condition) x 4 (syllable position) repeated-measures ANOVA was conducted using the ezANOVA function in R (Lawrence, 2022). Post-hoc comparisons were performed using paired-sample t-tests and emmeans (Lenth, 2025). The Bonferroni method corrected for multiple comparisons with the alpha level set at 0.05.

### H. Pupillometry

#### 1. Pre-processing

Pupil dilation from the left eye was recorded with the EyeLink 1000 at a sampling rate of 1000 Hz. Instances where participants had full or partial eye closure (e.g., blinks) were treated as missing data. Participants with excessive missing data (>50%) were excluded from further analysis. A total of 9 participants were excluded for this reason (refer to the Participants section in Methods).

Artifacts such as eye blinks were removed and replaced by linearly interpolating 50 ms before and 150 ms after an eye blink. A 4th order zero-phase, lowpass Butterworth filter at a 10 Hz cutoff was applied. Data were then epoched from -1 second to 7 seconds. To separate out tonic from phasic pupillary responses and account for between-participant differences in absolute pupil size, each trial was baseline corrected using a pre-cue interval (-0.5 to 0 sec). Specifically, the mean pupil size over the baseline window was computed for each trial. Tonic responses were calculated by averaging these baseline values across all trials; the across-trial standard deviation of these baseline values was also computed per subject. Phasic responses were calculated as z-scores based on the distribution of across-trial baseline responses. First, the trial-specific baseline was subtracted from the corresponding epoch. The resulting baseline-corrected time traces were then averaged and divided by the participant-specific baseline standard deviation to generate z scores. For robustness, we also computed phasic responses using two alternative approaches: baseline subtraction alone (no scaling) and proportional change relative to the trial baseline mean (data provided in supplemental figures FIGS S3A and S3B, respectively). These alternative analyses yielded similar results and supported the same conclusions; therefore, here we report z-scored pupil responses.

#### 2. Analyses

Tonic responses were compared across the different block types within each experiment (i.e., comparing anechoic and reverberant conditions in the main experiment and between uninterrupted and interrupted conditions in the follow-up experiment).

Three analyses were performed to quantify the effect of interruption and acoustic environment on the phasic pupil trace (described in further detail below): (1) cluster-based permutation testing on the entire pupil trace, (2) analysis of peak pupil dilation latency (time to maximum peak in seconds), and (3) analysis of peak pupil dilation amplitude. Peak pupil latency was computed as the time it took for the pupil dilation to reach its maximum peak. Peak amplitude was found by averaging pupil dilation across a ± 50 ms window around the peak latency.

To assess time-resolved differences in pupil dilation between interrupted and uninterrupted trials within each acoustic environment, we conducted a cluster-based permutation test on the baseline-corrected pupil time courses using custom MATLAB scripts based on the procedure described by (Maris and Oostenveld, 2007). For each acoustic environment (anechoic and reverberant), we compared the uninterrupted and interrupted conditions using a paired-sample (dependent) cluster-based permutation test across the entire pupil trace. We used 10,000 permutations with a two-sided threshold of p = 0.05 to define clusters of continuous time points exhibiting significant differences. The test statistic was the sum of t-values within each cluster (T-sum). The null distribution was generated by randomly sign-flipping the difference waveforms for each subject and constructing the same test statistic. Empirical clusters were considered significant if their T-sum exceeded the 95th percentile of the null distribution. This procedure controls for multiple comparisons across time while maintaining sensitivity to temporally sustained effects.

To examine the effects of reverberation and interruption on the latency and amplitude of the peak pupil dilation, peak pupil latency and amplitude were extracted for each participant and condition. Normality of the residuals was assessed using Shapiro-Wilk tests, which revealed that the data were not normally distributed (all p <0.05). Therefore, nonparametric statistical analyses were performed. For each participant, condition effects were quantified using subject-level difference scores. Main effects of interruption and environment were tested by sign-flipping these difference scores across participants to generate null distributions of the mean effect (10,000 permutations). Interaction effects were calculated by computing difference-of-differences for each participant and applying the same sign-flipping procedure. Two-tailed p-values were computed as the proportion of permuted effects whose absolute value exceeded the observed effect.

## III. RESULTS

### A. Behavioral Results

FIG 2 plots participants’ raw percent correct performance with and without the interrupter as a function of syllable position for each of the four tested conditions (uninterrupted/interrupted x anechoic/reverberant). For each syllable in both uninterrupted and interrupted conditions, performance was worse in reverberation than in the corresponding anechoic condition. Similarly, for every syllable and in both anechoic and reverberant conditions, performance was worse for interrupted compared to uninterrupted trials; however, the size of this interrupter cost was small for the syllable 1, before the interrupter, and greatest for syllable 2, right after the interrupter. Finally performance generally was best for the first syllable and second best for the last syllable, demonstrating well-known primacy and recency effects common in serial-recall tasks (Murdock, 1962).

**FIG. 2.**
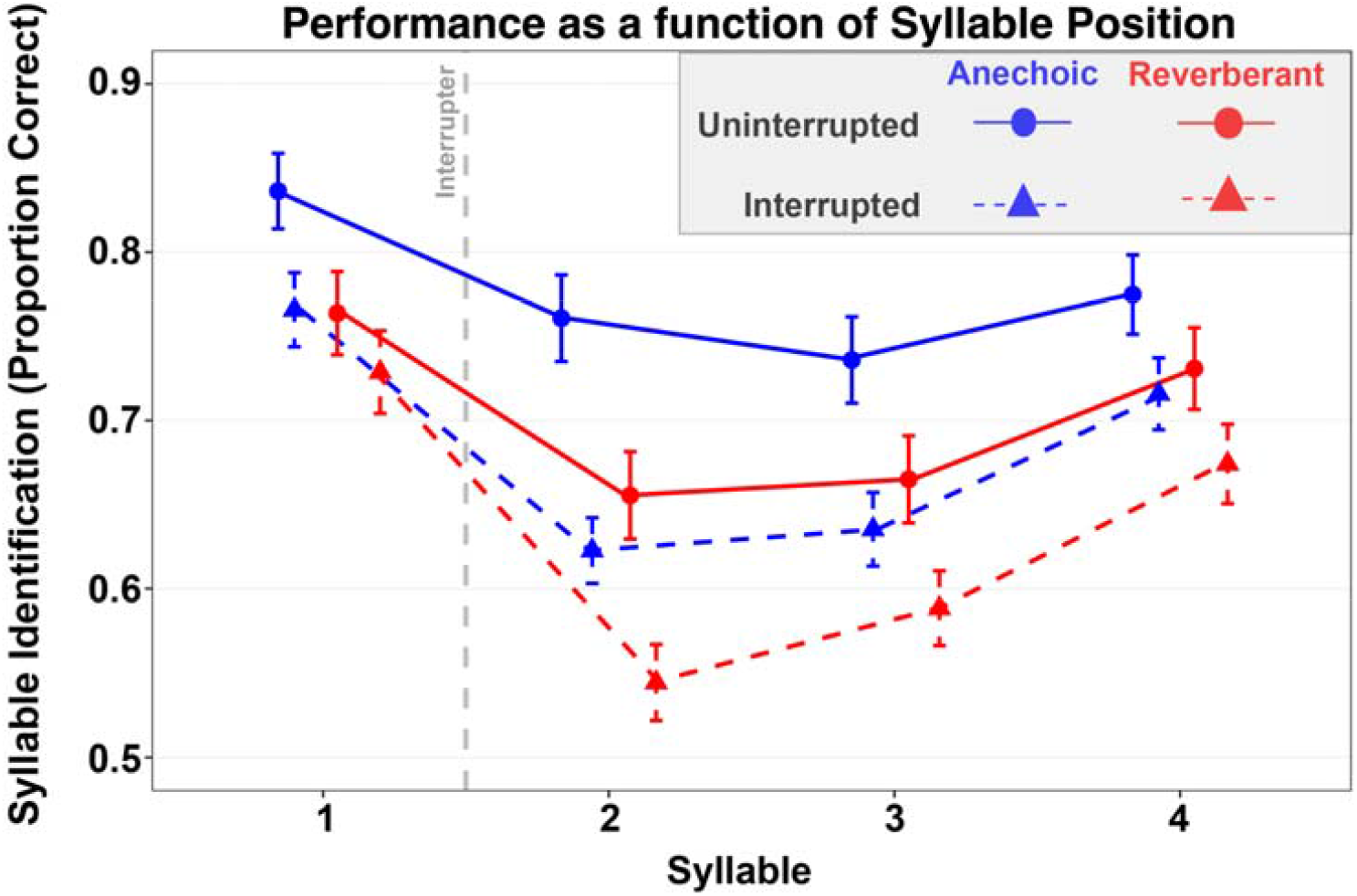
Mean syllable recall accuracy (Proportion correct) across syllable positions for uninterrupted and interrupted trials. Raw syllable recall accuracy across 4 conditions (anechoic/uninterrupted, anechoic/interrupted, reverb/uninterrupted, reverb/interrupted). Accuracy showed robust primacy and recency effects, with reduced performance for syllables 2–3. Both interruption and reverberation impaired performance, with an interaction trend suggesting larger interruption costs in anechoic trials. The error bars depict standard error of the mean.

These observations were supported by ANOVA results, which revealed significant effects of all three main variables of syllable position (F(3,177) = 95.03, p < 0.001), interruption condition (F(1,59) = 121.63, p < 0.001), and room environment (F(1,59) = 87.99, p < 0.001). Additionally, there were significant two-way interactions between syllable position and interrupted condition (F(3,177) = 16.38, p < 0.001) and between syllable position and room condition (F(3,177) = 7.69, p < 0.001), as well as a marginal interaction between interruption and room condition (p = 0.067). The three-way interaction between syllable position, interruption condition, and room environment was not significant (F(3,177) = 1.02, p = 0.386).

Overall, the results of the main ANOVA confirmed that subjects generally performed worse in reverberant than in anechoic conditions and worse in interrupted conditions than in uninterrupted conditions. To understand the various interactions between syllable position, room condition, and interruption, we conducted a follow-on analysis of the interruption effect—the size of the decrease in performance caused by the interrupter—for anechoic and reverberant settings as a function of syllable position.

FIG 3 plots the interruption effect as a function of syllable position for both anechoic and reverberant room simulations. The size of the interruption effect varied with syllable position. In particular, in both room simulations the interruption effect was largest for syllable 2, next largest for syllable 3, and smallest for syllable 1. In addition, the interruption effect tended to be larger in anechoic than in reverberant settings.

**FIG. 3.**
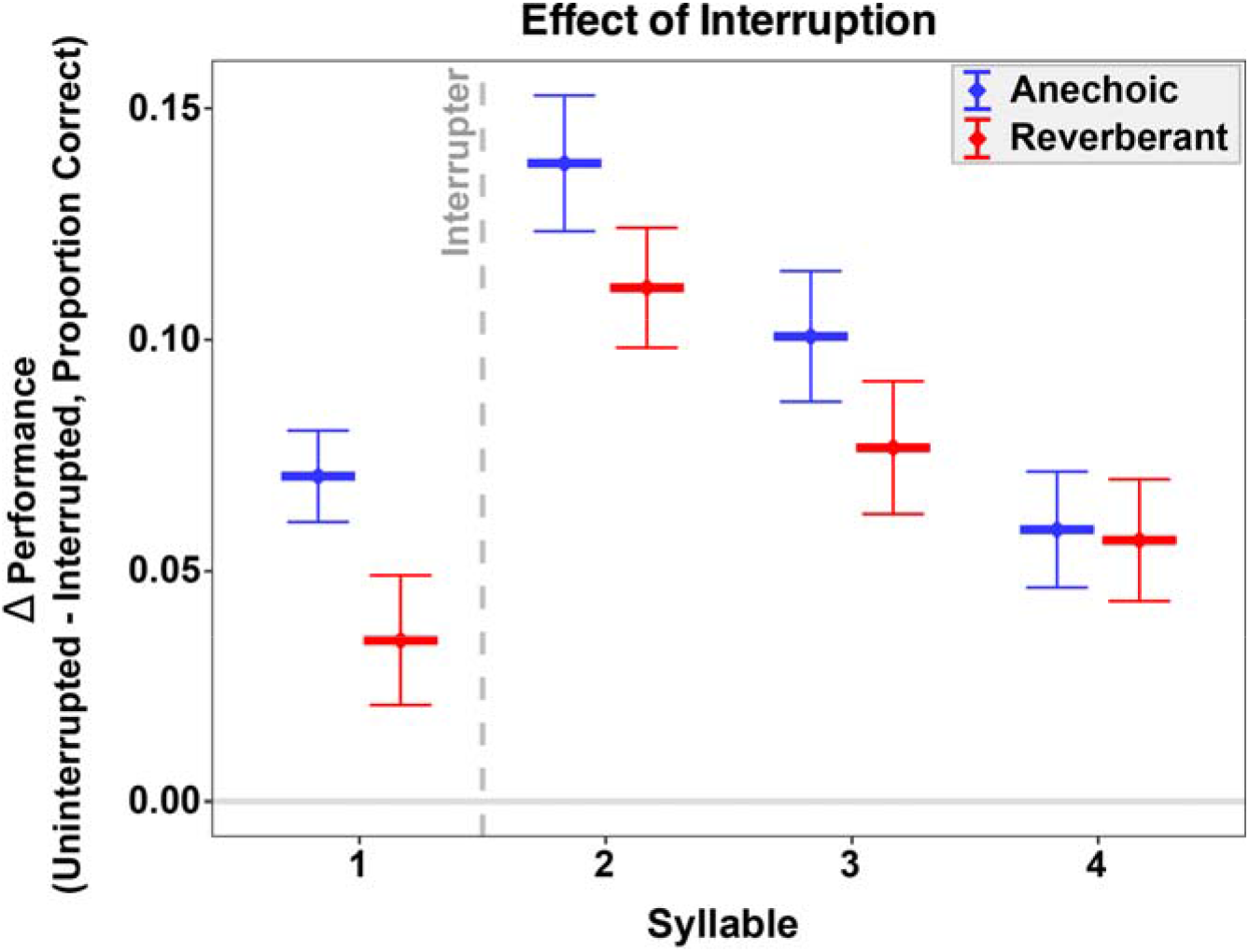
Effect of interruption on syllable recall accuracy across experiments. The effect of interruption was computed by finding the within-subject difference in syllable recall accuracy between uninterrupted and interrupted trials in the anechoic (blue) and reverberant (red) conditions. Error bars show the across-subject standard error of the mean. The grey dotted line depicts when the interruption occurred (before the onset of the second syllable)

The follow-up ANOVA on the interruption effect (defined as the difference in accuracy between uninterrupted and interrupted conditions), which included within-subject factors of room condition and syllable position, confirmed these observations. There was a significant effect of syllable position (F(3,177) = 16.29, p < 0.001). There was also a marginal effect of room condition (F(1,59) = 3.47, p = 0.067). Finally, the interaction between room condition and syllable position was not significant (F(3,177) = 1.02, p = 0.386).

Post-hoc comparisons showed that the effect of the interrupter was larger for syllable 2 than syllable 1 (Δ = 0.0719, p < 0.001), syllable 3 (Δ = 0.036, p = 0.042), and syllable 4 (Δ = 0.0669, p < 0.001). The effect of the interruption was also significantly larger for syllable 3 than syllable 1 (Δ = 0.0359, p = 0.0459) and syllable 4 (Δ = 0.0308, p = 0.0201), whereas syllables 1 and 4 did not differ (p=1.00). Post-hoc comparisons of the interrupter effect between anechoic and reverberant conditions showed a trend for larger effects in anechoic than reverberant when averaged across syllables (Δ = 0.022, p = 0.067).

### B. Pupil Dilation Results

#### 1. Tonic Responses

One might expect that in a testing block where stimuli were *consistently* more difficult than stimuli in another block (e.g. because the stimuli in a block were all in reverberation), there might be elevated pupil size based on increased arousal or anxiety. However, there were no significant differences in the tonic (i.e. uncorrected baseline pupil sizes across trials before the stimuli began) responses for anechoic and reverberant conditions in the main experiment (t(59) = 0.634; p = 0.5283), and there was no significant correlation between tonic response with behavioral performance across subjects. Similarly, in a follow-up experiment where the stimulus attributes were blocked by interruption rather than reverberation, the tonic response for interrupted and uninterrupted trials still did not differ significantly (t(14)=-1.717; p = 0.1079).

#### 2. Phasic Responses

In both anechoic and reverberant settings, trials evoked strong phasic pupil responses (FIG 4). In both settings, the interrupter caused a greater peak response in uninterrupted trials than in interrupted trials.

**FIG. 4.**
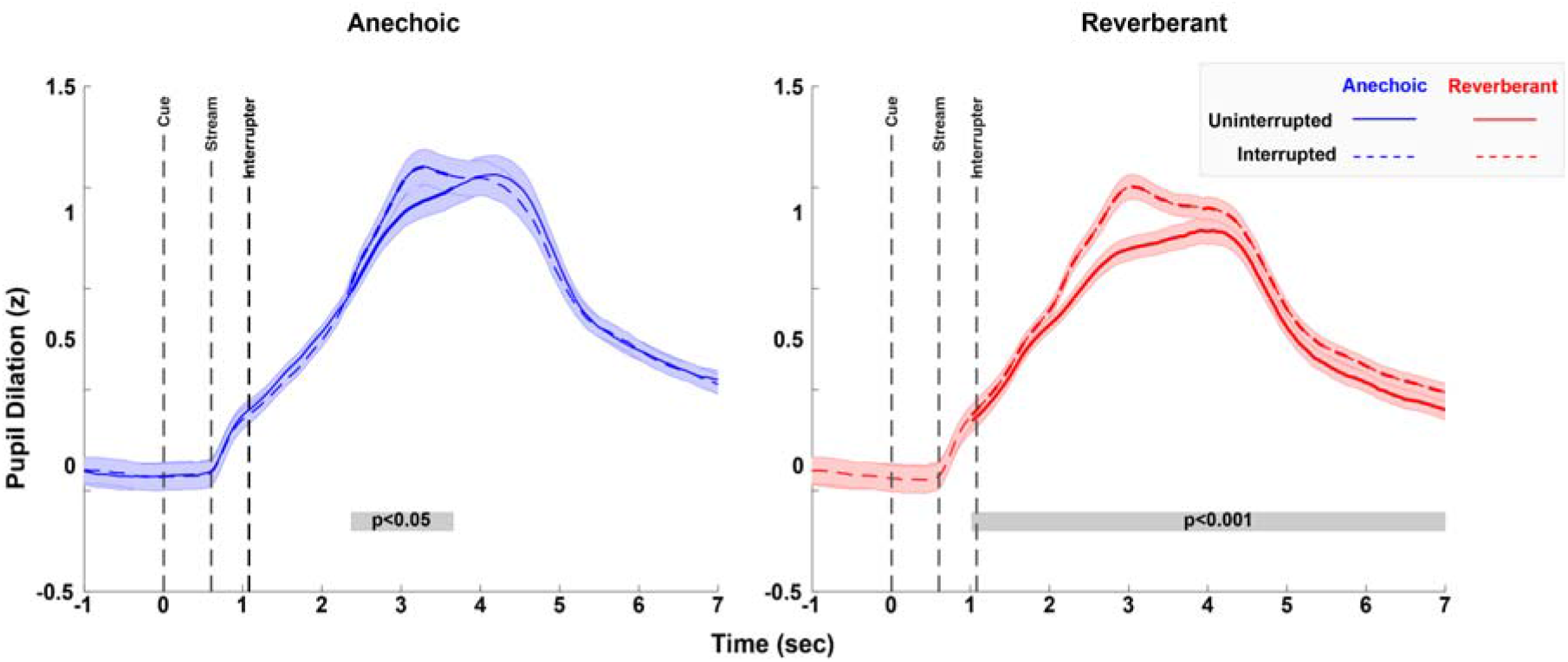
Across-subject average of pupil dilation time traces in the anechoic (left) and reverberant (right) conditions during uninterrupted (solid line) and interrupted (dashed lines) trials. Pupil traces are z-scored relative to baseline, based on the across-all-trial mean and standard deviation of the pupil diameter calculated over the baseline period 500ms before the cue onset (per subject). Error bars represent the standard error of the mean, computed across the subject means. The grey shaded line segments at the bottom of the two panels indicate time clusters that differed significantly between uninterrupted and interrupted conditions.

To test this statistically, we performed cluster-based permutation testing to identify the stretches of time where the phasic responses were greater for the interrupted condition. Significant clusters emerged in both anechoic and reverberant environments, indicating reliably greater pupil dilation in interrupted compared to uninterrupted trials. In the anechoic condition, a significant cluster was observed from approximately 2.31 to 3.81 seconds post-cue onset (p < 0.05), corresponding to the interval of peak pupil response. Similarly, in the reverberant condition, a significant cluster was observed over a longer stretch of time, from approximately 1.586 to 7 seconds post-cue onset (p < 0.001).

Peak pupil latency was computed by identifying the time of the maximum dilation for each subject’s aggregated data for each condition (FIG 5A). Permutation tests revealed earlier peak pupil latency in interrupted compared to uninterrupted trials (p < 0.001), and earlier in reverberant compared to than anechoic environments (p = 0.0026). The interaction between interruption and room condition was not significant (p = 0.72), indicating that the shift in peak latency due to interruption was comparable across acoustic environments.

**FIG. 5.**
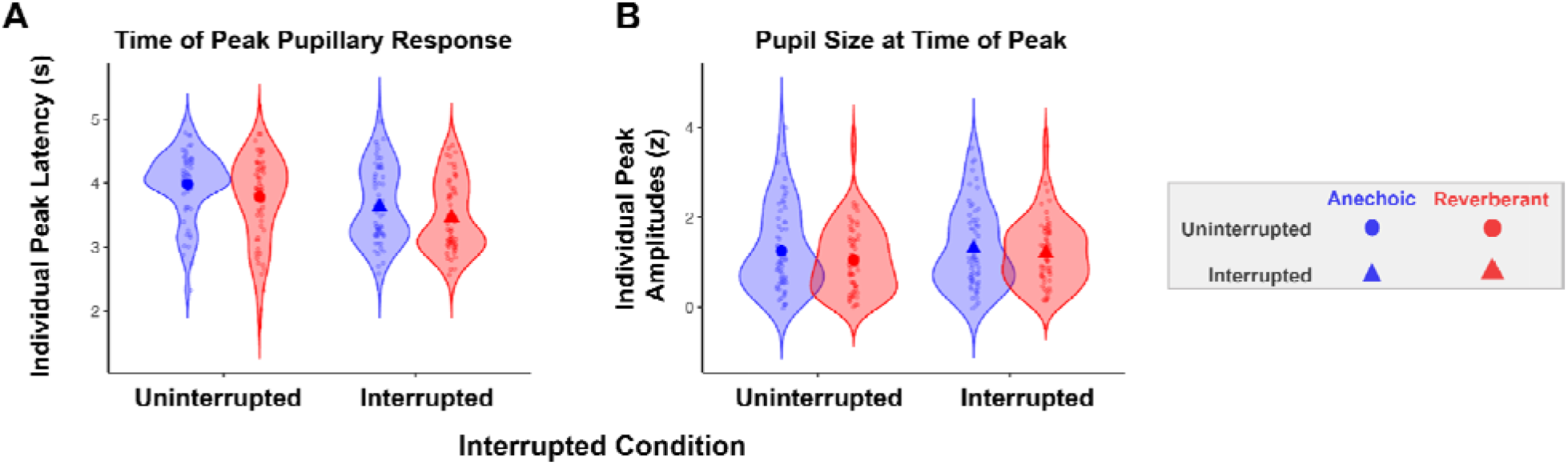
Pupil latency and amplitude across conditions. (A) Distribution of individual peak latencies and (B) distribution of individual peak amplitudes (z) as a function of acoustic environment and interruption, shown as violin plots with individual observations overlaid. Circle markers indicate uninterrupted conditions, triangle markers indicate interrupted conditions, blue indicates anechoic conditions, red indicates reverberant conditions.

As expected, peak pupil dilation was greater in interrupted trials than uninterrupted trials (p = 0.0001); however, it was unexpectedly greater in anechoic than in reverberant settings (p = 0.025) (FIG 5B). There was also a significant interaction between interruption and room condition (p=0.0018), driven by a larger difference between anechoic and reverberant peak pupil amplitudes for uninterrupted trials than for interrupted trials (uninterrupted trials Δ = 0.20 z-units vs. interrupted Δ = 0.10 z-units). That is, the effect of reverberation for trials *without* interruption was greater than for trials with interruption, indicating sub-additive effects of these two factors (each or which increases task difficulty). This interaction can be seen in the average pupil dilation traces in FIG 4, where the uninterrupted reverberant trials (solid red line) elicit a smaller phasic pupil dilation compared to the other conditions.

## IV. DISCUSSION

This study examined how auditory spatial attention, stimulus interruption, and reverberation jointly shape auditory perception and listening effort. Using a selective attention task with interleaved, spatially separated syllable streams with and without interruptions in anechoic and reverberant environments. In addition to behavioral performance, we simultaneously measured pupil dilation to provide a complementary index of attentional allocation and cognitive load. We considered two competing behavioral hypotheses: 1) Reverberation would smear the acoustic features of the mixture, reducing the salience of the interrupter and decreasing its disruptive impact, or 2) reverberation would make the task more difficult overall, compounding the effect of the interrupter. Physiologically, we hypothesized that reverberation would increase task difficulty generally, which would be reflected in larger pupil dilation, particularly in the tonic component of the pupil response, relative to anechoic conditions.

### A. Effects of Reverberation on Behavioral Performance

Many of our behavioral results replicated findings from the previous experiments using variations of the current experimental paradigm (Figarola et al., 2025; Liang et al., 2022, 2025). Specifically, we found performance tended to be best for the first syllable and second best for the final syllable, in line with classic serial-recall phenomena in which early (primacy) and late (recency) list items are remembered best (Murdock, 1962). Also consistent with past, similar studies, the interrupter significantly impaired syllable recall performance in both anechoic and reverberant trials (Figarola et al., 2025). In the current study, specifically, interruption impaired syllable recall accuracy for syllables 1-3, and, as in past studies, the largest effect was on recall of the syllable immediately following the disruption (Syllable 2). This pattern is consistent with prior work on attentional capture, in which an abrupt stimulus disrupts selective attention and interferes with ongoing stream segregation and memory encoding (Figarola et al., 2025; Liang et al., 2022; Luthra, 2024; Mondor and Zatorre, 1995; Parmentier, 2014; Parmentier et al., 2011). This disruption can be especially costly when the interruption is temporally close and overlaps with the subsequent syllable, where limited time is available to reestablish attentional focus (Larson and Lee, 2013; Logan, 2005). However, participants were able to recover towards the end of a trial, demonstrating that they were able to refocus attention on the target stream. These dynamics highlight the everyday demands made on attentional control in complex auditory environments.

While many of the current behavioral results align with past findings, there was one interesting difference: The effect of reverberation on performance in the current experiment was much larger and more consistent than what was reported previously in a very similar study (Experiment 4 in (Figarola et al., 2025). This previous study used the same task, the same anechoic and reverberant spatialization methods and rooms, the same syllable set, the same blocking of trials, and the same participant interface. In the current experiment, performance was consistently worse in reverberant than in anechoic trials (ranging between about 5%-10% difference in percent correct performance) both for uninterrupted and interrupted trials. Yet in the very similar previous study, anechoic and reverberant performance was nearly identical across the two room conditions for all syllables in the uninterrupted trials and for all syllables except the one occurring right after the interruption in interrupted trials.

The most notable methodological differences between the two experiments were that the current study:

- was conducted in a laboratory setting, rather than online, in order to allow us to track pupil dilation over the course of each trial;
- added extra time in between trials to allow phasic pupil responses to return to baseline after each trial; and
- used four syllables in each target and masker stream, rather than five, to reduce the duration of individual trials.

We believe that the key difference lies in the difference in the length of the target and distractor streams. In the current experiment, with only four syllables, the working memory load is lower, which places greater emphasis on perceptual differences related to the room acoustics. This in turn may lead to more consistent effects of reverberation on performance: When cognitive demands on working memory determine performance, outcomes may be less tightly coupled to the room acoustics. However, the fact that the prior study was conducted online may also play a role, leading to differences in everything from the degree of experimental control to the demographics and motivation of the participants. Future work could explore these factors to confirm our interpretation that reducing the memory load accounts for why reverberation has a large impact on performance in the current experiment, even though it had a limited impact in the previous, very similar study.

Finally, in the current study, the interrupter tended to have a larger effect in anechoic than in reverberant trials. We believe this reflects effects of reverberation on the perceptual salience of the interrupter. Late reverberation from the preceding target and distractor syllables overlaps with the interrupter, energetically masking its onset (Darwin and Hukin, 2000; Nabelek et al., 1989; Ruggles and Shinn-Cunningham, 2011; Shinn-Cunningham, 2000). Reverberation also reduces the clarity of the scene by blurring segregation cues, such as spatial and pitch differences (Darwin and Hukin, 2000; Devore and Shinn-Cunningham, 2003; Nabelek et al., 1989; Shinn-Cunningham et al., 2005). In contrast, in anechoic environments these details remain better defined, so the interrupter may be more salient and more disruptive. Thus, though the interrupter impaired performance in both environments, its greater impact in anechoic conditions suggests that attentional capture from an unexpected event is stronger when there is greater perceptual clarity of the interrupting event.

### B. Effects of Reverberation on Pupil Responses

As noted above, performance was significantly worse in reverberant blocks than anechoic blocks. Importantly, reverberant performance still exceeded chance performance, demonstrating that participants were actively engaged in the task even in these difficult, reverberant trials (i.e. the difficulty of the condition did not cause them to give up). We hypothesized that participants would have stronger pupillary responses in reverberant blocks compared to anechoic blocks, reflecting their increased difficulty; pupil dilation is often argued to increase with task difficulty and cognitive load, whether from acoustic degradation, linguistic complexity, or memory demands (Contadini-Wright et al., 2023; Koelewijn et al., 2012, 2015; Winn et al., 2015). However, the current results suggest that attentional salience strongly drives the pupillary response in a way that was not explicitly tested by these previous studies.

Tonic (baseline) pupillary responses were no different across reverberant and anechoic blocks. That is, reverberation did not substantially elevate *sustained* arousal or global listening effort in this paradigm, even though reverberation diminished behavioral performance. Additionally, there was no tonic pupil size difference across blocks defined by the presence or absence of interruption (see supplemental follow-up study). Together, these observations suggest that if sustained effort is elevated by the expectation of ongoing stimulus difficulty, the stimulus conditions in the current experiment did not rise to the level of eliciting that sustained difference.

Even more surprising, peak pupil dilation was significantly larger and later in anechoic trials compared to reverberant trials. This further reinforces the notion that performance accuracy on its own – which was higher in the reverberant condition – cannot explain the magnitude of phasic pupil responses. A similar mismatch between performance and elicited pupil size has previously been observed for sentence recognition tasks as well (Winn and Teece, 2021, 2022).

One possible explanation for the reduced phasic pupil responses in reverberation is that participants did not engage as fully on the task in the more difficult, reverberant-trial blocks. For instance, the FUEL framework proposes that effort depends not just on task demands but also on task engagement, motivation, and the perceived likelihood of success (Pichora-Fuller et al., 2016). Following this, studies have reported larger pupil dilation for acoustically clearer or more informative speech, suggesting that pupil responses track a participant’s willingness to invest effort when it is likely to yield a benefit (Fitzgerald et al., 2024; Mechtenberg et al., 2024). Although task performance was not in the sub-40% range that some studies have identified as resulting in effort withdrawal (Ohlenforst et al., 2017; Wendt et al., 2018), listeners still may have adapted to the reverberant condition, neutralizing the novelty of this particular challenge. To directly test this idea, we conducted a follow-up study in which we varied the blocking of trials so that harder reverberant trials were randomly intermingled with easier anechoic trials, so that listeners could not predict whether an upcoming trial was difficult or easy. Specifically, over the experiment the same distribution of trials was presented, but half of the blocks contained only uninterrupted trials and half contained only interrupted trials (Supplementary FIG S1 and S2). Phasic responses were generally lower in this supplemental experiment, likely reflecting the fact that the structure of the stimuli within a block were identical, and therefore less surprising overall. Critically, just as in the main experiment, for uninterrupted trials the phasic response was lower for reverberant than for anechoic simulations—even though the two trial types were intermingled. This makes it less likely that differences in the phasic responses for uninterrupted trials reflect differences in participant effort, engagement, or long-term expectations over an entire testing block.

We argue that reverberation results in weaker phasic pupil dilation because it degrades segregation cues, making the onsets of individual syllables less distinct and thus less attentionally salient. Specifically, in anechoic trials, each successive syllable may have evoked a phasic response that added to the accumulated pupil dilation over the course of the trial, leading to larger peak responses that built up over a longer time. In reverberant trials, each syllable is less clear, driving a weaker phasic response. In this view, rather than reflecting task difficulty, phasic responses index the acoustic salience of individual syllable onsets (see Klingner et al. 2011 and Winn et al. 2018 for further examples of the buildup of individual components to a larger pupil response).

In both acoustic environments, the interrupter elicited a large, robust pupil response, consistent with salient and behaviorally relevant events eliciting pupil dilation (Dayan and Yu, 2006). The cluster-based permutation analyses confirmed sustained increases in pupil dilation for interrupted relative to uninterrupted trials in both room types. Moreover, interruption shifted peak latency earlier compared to uninterrupted trials. These results confirm that in both anechoic and reverberant trials, the interrupter was salient, driving a robust phasic pupil response, consistent with rapid orienting to a salient, unexpected event. Given that the target and distractor streams were speech syllables dominated by a consistent vowel sound that would carry strongly in reverberation, but the interrupters were brief, non-speech sounds, this result makes sense—the reverberation can cause syllables spoken by the same talker to smear together and be difficult to segregate, but do not disrupt segregation of the novel interrupters, which are acoustically distinct from the target and distractor streams. Consistent with this, the difference in peak pupil dilation between anechoic and reverberant trials was smaller in interrupted trials than in uninterrupted trials. This conforms to the idea that the early interrupter-evoked response, which dominated the phasic pupillary responses, had a similar strength in both rooms.

The strong effect of the interrupter is consistent with pupil dynamics tracking the timing of attentional shifts, cognitive load, and inhibitory control (Koelewijn et al., 2014, 2015; McCloy et al., 2017; Winn and Moore, 2018; Zekveld et al., 2018). Recent work further suggests that this phasic pupil response reflects activity of the locus-coeruleus-norepinephrine (LC-NE) system, which plays a key role in processing interrupting events in the environment (Dayan and Yu, 2006; Lim et al., 2021; Sara and Bouret, 2012). Through this mechanism, the pupil-linked LC-NE system is particularly sensitive to abrupt, bottom-up changes in the auditory scene, (Zhao et al., 2019a), making it a reliable marker of disrupted attention. In addition, the interrupter increases cognitive demands by requiring participants to refocus attention back on the target stream after the involuntary disruption. The compounding effects of the interrupter align with Load Theory, which proposes that interference from irrelevant stimuli increases when perceptual and cognitive resources are taxed (Lavie et al., 2004). In the present study, the phasic response triggered by the interrupter may thus reflect a combination of both a salience-driven attentional reset and an increase in cognitive load, with the enhanced pupil response indexing this compounded burden.

Taken together, the pupil data suggest that pupil dilation in this task is driven more strongly by discrete, salient acoustic events and dynamic processing demands than by sustained cognitive effort related to performance accuracy. The spectrotemporal smearing caused by reverberation degrades performance because it makes it difficult to segregate the competing target and masker syllables, which are identical except in their direction. However, the interruptions—abrupt and behaviorally salient events that are distinct in their spectrotemporal properties from the speech—elicits large, rapid pupil responses in both anechoic and reverberant environments. Thus, pupil dynamics reveal changes in sensory scene organization and salience that are not directly inferred from behavioral accuracy alone.

### C. Future Directions

Future work should explore whether similar salience-driven pupil dynamics dominate responses for more naturalistic and linguistically rich materials. Dominance of acoustic salience was observed by Winn and Teece (2021) in sentence recognition tasks where a disruption was either a clear departure from the ongoing speech stream (a noise masking a word) versus a change within the speech stream itself (a mispronunciation of a word in a sentence). In their study and in the current study, the disruptor that was more obviously different from the target stream elicited stronger pupil dilations. However, this previous work was not designed with salience explicitly as a stimulus factor, leaving opportunity to examine this issue in greater depth. Considering the greater pupil dilation for vocoded (i.e. less perceptually salient) speech versus clear speech (Winn 2016), there is likely a competing tension between pupil dilation driven by effort to complete a task versus dilation driven by stimulus salience. Following up on the current work, manipulating reverberation strength parametrically would allow characterization of the relationship between acoustic smearing and phasic response magnitude. In addition, systematically varying working memory load within the same laboratory paradigm would directly test the proposal that reducing working memory demands increases the relative contribution of acoustic factors to performance.

Expanding the participant population is also important to ensure that the current results are not over generalized. Older adults, listeners with hearing impairment, and individuals with attentional or cognitive differences may show distinct interactions between reverberation, interruption, and pupil-linked indices of effort. Individual differences in working memory capacity and attentional control could alter how reverberation and bottom-up interruptions interact when participants deploy top-down selective attention.

Finally, combining pupillometry with electrophysiological measures (e.g., EEG or MEG) could tease apart the contributions of early sensory encoding, attentional reorienting, and memory-related processes during interrupted attention. Such multimodal approaches could clarify how cortical dynamics interact with pupil-linked neuromodulatory signals during sustained auditory attention in complex environments.

### D. Conclusions

Reverberation degrades acoustic scene clarity, reducing the distinctiveness of temporally adjacent events and impairing segregation of competing speech streams. In the present task, this degradation produced reliable declines in syllable recall performance. Critically, reverberation also reduced phasic pupil responses during uninterrupted trials, indicating that individual syllables were less salient when temporally smeared by room reflections. These physiological results provide converging evidence that reverberation weakens event-level sensory encoding rather than simply increasing global cognitive effort.

Despite the reduction in clarity caused by reverberation, abrupt interruptions with clear spectral deviations from the ongoing syllable stream elicited robust behavioral disruption and large phasic pupil responses in both anechoic and reverberant environments. Peak pupil responses occurred earlier and were significantly larger in interrupted trials, consistent with a framework of pupil dilation driven by rapid attentional capture by salient, unexpected events. Importantly, the interrupter-evoked pupil response was comparable across room conditions, suggesting that perceptually distinct, novel sounds retain their salience even in strongly reverberant environments.

Together, these findings demonstrate that pupil dynamics in this paradigm are driven more strongly by discrete, bottom-up acoustic salience than by sustained task difficulty. Although reverberation impaired performance, it did not elevate tonic pupil diameter and instead reduced phasic responses to ongoing speech events. In contrast, sudden interruptions dominated the pupillary response regardless of room acoustics.

These results highlight a dissociation between behavioral difficulty and pupil-linked arousal in complex listening environments and underscore the value of pupillometry for revealing dynamic changes in attentional allocation that are not apparent from accuracy measures alone.

## SUPPLEMENTARY MATERIAL

See supplementary material for follow-up behavioral and pupil dilation results (S1-S2) as well as baseline correction methods (S3).

## ACKNOWLEDGMENTS

Research reported in this publication was supported by the National Institute On Deafness And Other Communication Disorders of the National Institutes of Health (T32DC011499 to VF, R21- DC018408 to CAB, R01-DC019126 to BGSC, and R01-DC022699 to BGSC & CAB) and the Office of Naval Research (N00014-19-12332, N00014-20-12709, and N00014-23-12065 to BGSC). SL was supported by NIH NRSA F32DC020625. We would also like to thank Grace Swatsworth for her help in data collection.

## AUTHOR DECLARATIONS

### Ethics Approval

This study was reviewed and approved by Carnegie Mellon University Institutional Review Board. The participants provided their formal written consent to participate prior to their participation in this study.

## DATA AVAILABILITY

The raw data supporting the conclusions of this article will be made available by the authors, without undue reservation.

