## Supplemental Material for "Acoustic Salience Drives Pupillary Dynamics in an Interrupted, Reverberant Task"

**Supplementary Material**

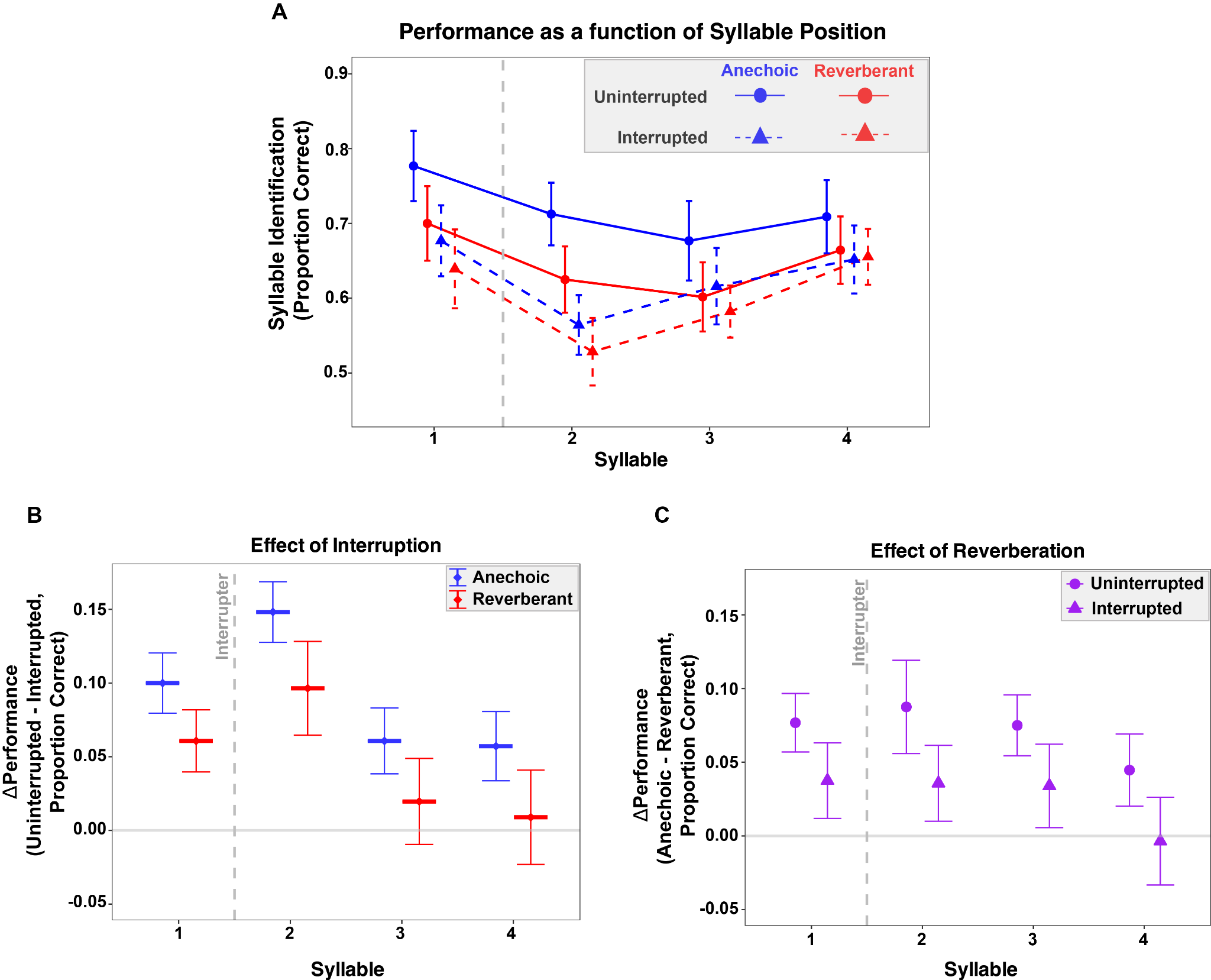

**FIG. S1.** **Behavioral Performance in the follow-up study replicates the main effects of interruption and reverberation.** Behavioral accuracy (Proportion correct) from the small-scale study (N=14; one participant not included due to missing behavioral data), plotted as a function of syllable position, interruption, and acoustic environment. Error bars represent across-subject standard error. (A) Accuracy for all four experimental conditions, showing a significant effect of syllable position (F(3,29)=5.29, p=0.0036). (B) Effect of interruption (uninterrupted - interrupted), averaged across the two acoustic environments. Interruption significantly reduced target syllable recall overall (F(1,13) = 23.35, p <0.001), with a larger cost at earlier syllables. (C) Effect of reverberation (anechoic - reverberant), averaged across interrupted conditions. Reverberation significantly impaired performance (F(1,13) = 12.85, p = 0.003), with the largest effects observed in the middle syllables. There was also a significant interaction between interruption and environment (F(1,13) = 5.22, p = 0.040).

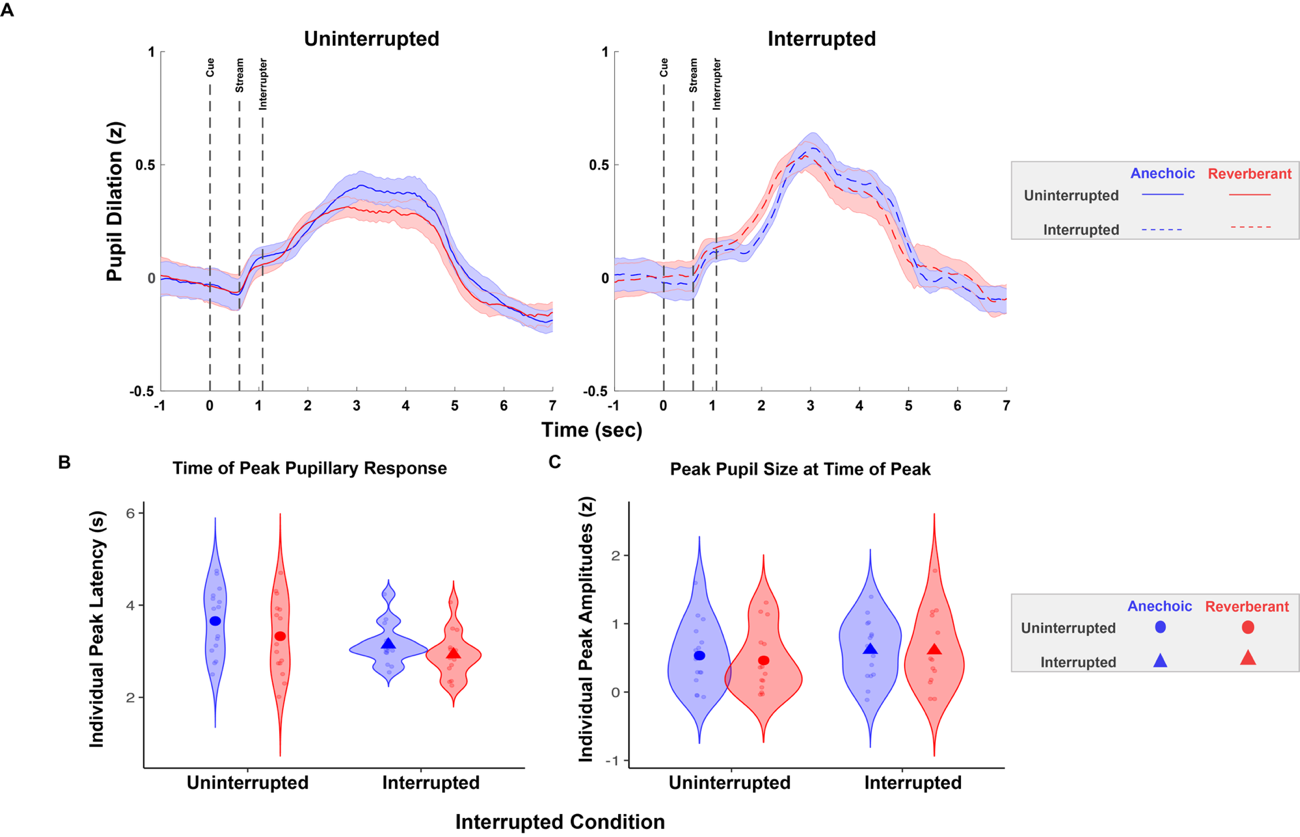

**FIG. S2. Follow-up study pupil responses blocked by interrupted condition (N=15).** (A) Pupil dilation time traces in the uninterrupted (left) and interrupted (right) conditions during anechoic (blue) and reverberant (blue) trials. These pupil traces are z-scored and baseline corrected (-500ms pre stimulus onset). The error bars represent the within-subject standard error. For (B) and (C), the y-axis are uninterrupted and interrupted conditions. Circle markers indicate uninterrupted conditions, triangle markers indicate interrupted conditions, blue is anechoic conditions, and red is reverberant conditions. (B) Time to peak (seconds) across four conditions. Permutation tests revealed that peak latency was significantly earlier for the interrupted than uninterrupted trials (p=0.005) and earlier in reverberant than anechoic environments (p=0.002), with no interaction (p=0.5828). (C) Peak pupil amplitude (z-scored) across conditions. Peak amplitude showed a significant effect of interruption (p=0.0051), with larger peaks in interrupted than uninterrupted trials, but no main effect of reverberation (p=0.413) and no interaction (p=0.2248).

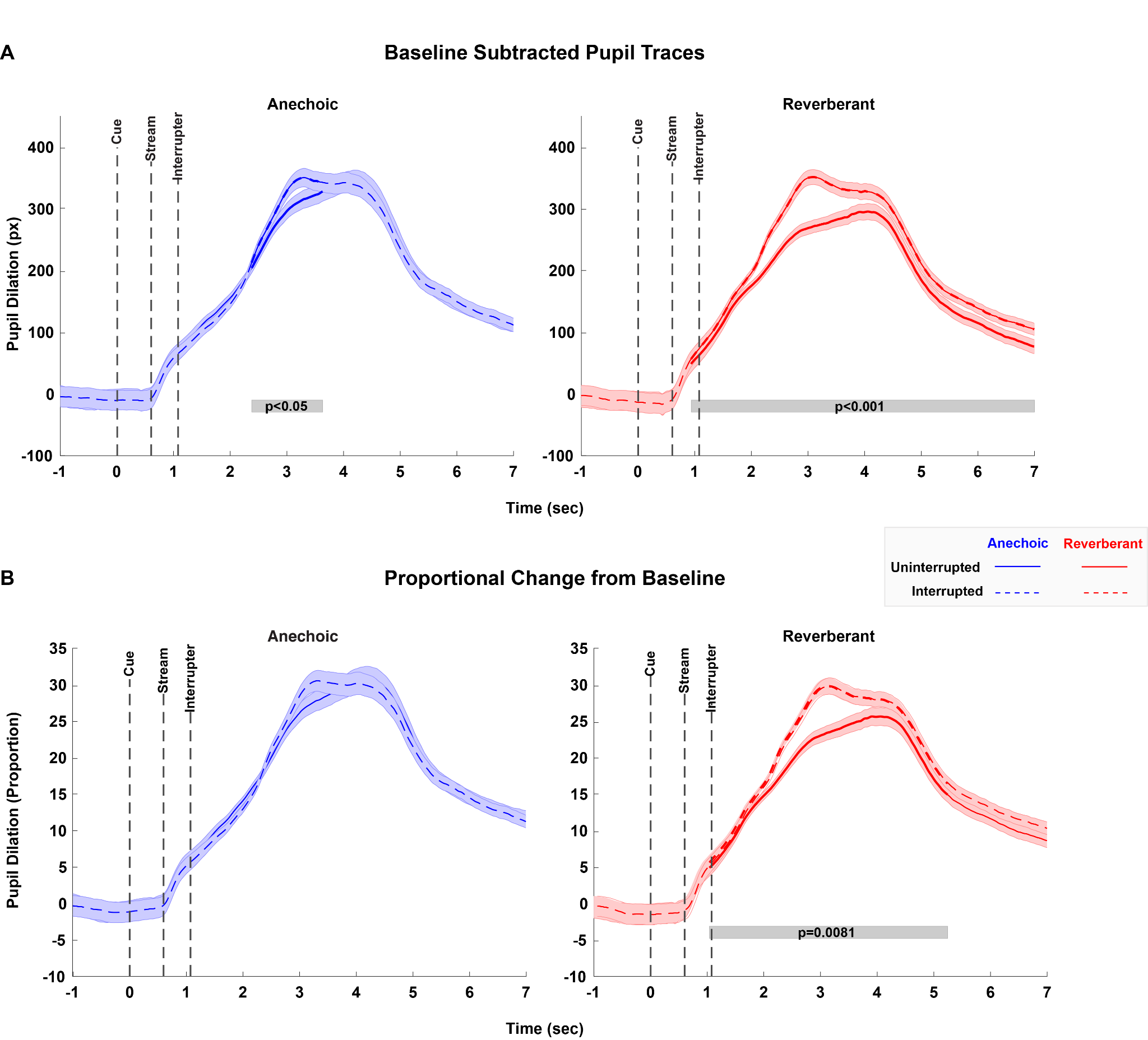

**FIG. S3.** (A) Baseline subtracted pupil traces. (B) Proportional Change from Baseline. For each panel, anechoic traces are in the left (blue) and reverberant traces are on the right (red), with uninterrupted (solid line) and interrupted (dashed line) trials. The error bars represent the within-subject SEM.
